# The Calcium-sensing receptor is an important regulator of female visceral and perivascular adipocyte function

**DOI:** 10.1101/2023.08.02.551589

**Authors:** Erin Higgins, Zaniah Gonzalez Galofre, John McAbney, James Leiper, Laura Dowsett

## Abstract

The Calcium-sensing receptor (CaSR) is a G protein-coupled receptor activated by fluctuations in extracellular calcium concentrations. Its importance in calcium homeostasis has long been established, though its role in other tissues is not well understood. Obesity is a major epidemic with both clinical and social consequences, therefore, understanding the full function and regulation of adipocytes is of critical importance. Adipocyte CaSR has previously been linked to lipolysis and inflammation *in vitro*. In this study, we set out to further our understanding of adipocyte CaSR *in vivo* via the generation of an adipocyte specific CaSR knockout mouse (CaSR^Ad-/-^). We found female CaSR^Ad-/-^ mice weighed less than wildtype littermates with a significant reduction in visceral adipocyte size potentially due to increased expression of brown fat markers UCP-1 and Cidea. We also established that CaSR is expressed in perivascular adipose tissue (PVAT) and that its deletion protects female mice from PVAT driven hypercontractility. In contrast CaSR deletion had no effect on male body mass, adipocyte size or vascular reactivity. In conclusion, CaSR seems to be of importance in female but not male visceral and perivascular adipose tissue.

## Introduction

The Calcium-sensing receptor (CaSR) is a key regulator of plasma calcium concentrations through its expression in the parathyroid gland, bone, and kidney [1-3]. However, beyond its role in calcium homeostasis, CaSR is found in many cell types and its function in this wider context is less well understood. Adipocyte expression of CaSR was first detected in primary adipocytes as well as pre-adipocytes isolated from human visceral adipose tissue [4]. This was subsequently confirmed in several adipocyte cell lines including LS14, SW872 and 3T3-L1s [5-7]. *In vitro* studies have shown CaSR to be anti-lipolytic through the suppression of cAMP signalling [6, 8], and to play a role in adipocyte inflammation via the stimulation of NF-κB signalling and the NLRP3 inflammasome [5, 9]. Given that the regulation of lipid accumulation and pro-inflammatory signalling are both considered significant pathways in the development of type 2 diabetes and the metabolic syndrome, these studies suggest that disrupted CaSR signalling would be detrimental to adipose health [10]. However, until recently all mechanistic studies examining the role of CaSR in adipocytes have been performed *in vitro* which limits the understanding of its role in whole body metabolism.

Early transgenic models demonstrated a clear role for CaSR in calcium homeostasis, with global CaSR deletion leading to severe hypercalcaemia, skeletal deformities and early death [3, 11]. As global CaSR deletion is lethal at a young age the Cre/loxP system was utilised to develop tissue specific CaSR transgenic mouse lines [1, 3]. By targeting exon 3 extensive disruption of both the CaSR transmembrane and Ca^2+^ binding domains occurs, and this model has been utilised to demonstrate the significant role of CaSR in the kidney [1]. A recent study by Sundararaman *et al*. [12] developed the first adipocyte specific CaSR knockout mouse; however, this mouse was further crossed to an ApoE^-/-^ mouse to investigate the role of CaSR in atherogenesis. Although Sundararaman *et al*. found no effect of CaSR deletion on body weight, triglyceride levels or adipose cytokine expression the absence of ApoE complicates the assessment of the role of CaSR in adipocyte homeostasis as ApoE deletion alone is known to reduce fat mass and adipocyte size [13]. Therefore, in this study we set out to establish whether CaSR deletion alone affects adipocyte function.

Currently, the function of CaSR has only been explored in models of visceral adipose tissue. Perivascular adipose tissue (PVAT) is a specialised adipose depot found around major conduit and resistance arteries. PVAT secretes a wide range of factors including adipokines, cytokines and vascular signalling molecules such as NO and AngII which all contribute to vascular reactivity [14]. CaSR expression within perivascular adipose tissue has not yet been investigated. Using our transgenic mouse model, we set out to establish whether CaSR in PVAT is a regulator of vascular reactivity. Given the known differences between male and female adipose physiology and distribution, we compared the effect of CaSR deletion between the sexes.

## Methods

### Mice

All animal procedures were conducted in accordance with the United Kingdom Home Office Animals (Scientific Procedures) Act, 1986, and directive 2010/63/EU of the European Parliament on the protection of animals used for scientific purposes. CaSR floxed mice (CaSR^fl/fl^) previously generated by Toka *et. al*. [1] and deposited in the Jackson Laboratory repository (stock #030647; Casr^tm1Mrpk^) were crossed with mice possessing the AdipoQ-Cre BAC transgene (stock #028020; B6.FVB-Tg (AdipoQ-Cre)1Evdr/1) to ensure CaSR expression was abolished in mature adipocytes only (CaSR^Ad-/-^). Jackson Laboratory protocols were followed for standard genotyping. To confirm successful deletion in primary adipocyte, primers surrounding exon 3 and the neomycin cassette (Supplementary Figure 1a and b, Primer sequences found in Supplementary Table 1) were used as described by Toka et al. Following weaning, male and female CaSR^fl/fl^ and CaSR^Ad-/-^ mice were weighed weekly until 18 weeks of age. Body weight is expressed as mean mass (g) ±SEM. Male and female C57Bl/6 mice were purchased from Envigo and used between 8-12 weeks of age.

### Free Fatty Acid

Free fatty acid (FFA) content was quantified in mouse plasma samples using the Free Fatty Acid Quantitation Kit (Sigma, #MAK0044) in line with the manufacturer’s instructions. Absorption was measured at 570nm using the SPECTRAmax 190 Microplate Spectrophotometer system (Molecular Devices, UK). Plasma FFA concentration is provided as mean concentration (nmol/μl) ±SEM.

### Triglycerides

Plasma triglyceride content was assessed in mice using the Triglyceride Assay Kit (Abcam, #65336) in line with the manufacturer’s instructions. Absorption was measured in duplicate at 570nm using the SPECTRAmax 190 Microplate Spectrophotometer system (Molecular Devices, UK). Plasma triglyceride concentration is provided as mean concentration (nmol/μl) ±SEM.

### Histology

Adipose tissue was excised, washed in PBS, and placed in ice-cold 4% paraformaldehyde (PFA) for 48h. Tissues were then placed in 70% ethanol for short-term storage. Tissue samples were dehydrated by the Shandon Excelsior ES system (ThermoFisher, UK) and embedded in paraffin blocks using the Shandon Histocentre 3 (ThermoFisher, UK). Paraffin blocks were sectioned at 5μm, mounted on silane-treated slides and cured at 60°C for 1h, then at 45°C for 12h. Following deparaffinization, slides were subjected to haematoxylin and eosin staining and mounted using DPX mountant (Sigma, #06522).

Image analysis was performed by assessing two tissue sections, separated by an interval of >30μm. Three images were collected from each section using the Olympus BX41 microscope system at x10 magnification. Cell size and number were measured manually (Image J: v1.52). In each image, 50 adjacent cells were measured and averaged to give mean cell area (AU) ±SEM in each condition. Cell number was assessed by counting the number of cells in a 500 pixel^2^ area, expressed as mean cell number ±SEM.

Aortas were excised, and PVAT was removed. Tissues were briefly washed in ice-cold PBS and fixed overnight in 4% PFA at 4°C. The following day, the aortas were dehydrated in 15% sucrose/PBS until they sank and then 30% sucrose/PBS. Aortas were cut in three sections and embedded in optimal cutting temperature (O.C.T) compound (Tissue-Tek), frozen in 100% ethanol cold vapours in dry ice and stored at -20°C until used. Sections 10μM thick were cut using an Epredia Cryostar NX50 cryostat and kept at -80°C until use. Slides were removed from the -80°C freezer and left wrapped in foil for 20min, then unwrapped and left for further 10min. Slides were fixed in ice-cold acetone/ethanol (75%/25%) for 10min, air dried for 10min and then placed in running water. Slides were stained with haematoxylin and eosin and dehydrated in 90% alcohol followed by 100% alcohol and xylene. Slides were then mounted using DPX mounting medium (Sigma) and imaged at 10x using an EVOS microscope.

The aortic wall thickness was measured in each sample at four different locations and the mean of these values was calculated. Between 8 to 12 sections per sample were measured and then averaged.

### Primary Adipocyte Isolation

Gonadal and inguinal adipose depots were dissected and placed immediately in ice-cold PBS. Adipose pads were finely minced and incubated in serum-free Optimem supplemented with 1mg/mL collagenase A (Sigma, #10103578001) for 1h at 37°C. Adipocytes were mechanically dissociated by pipette aspiration at 10min intervals. Adipose digests were centrifuged at 800rcf for 5min at 22°C. Digests were washed in PBS and the centrifugation step was repeated, causing fractional separation of adipocytes and stromal-vascular cells.

### qRT-PCR

Whole adipose tissue or isolated primary adipocytes and aorta were mechanically homogenised with 5mm stainless steel beads using a TissueLyser II system (Qiagen, UK) in the presence of 700μL QIAzol Lysis Reagent (Qiagen, #79306). RNA was isolated using standard phenol/chloroform separation, with the clear aqueous fraction taken forward for RNA clean-up using the RNeasy Mini Kit (Qiagen, #74106). cDNA synthesis was performed using the High-Capacity cDNA Reverse Transcription Kit (Invitrogen, #4368814), according to the manufacturer’s protocol. Primers (Supplementary Table 2) were used at 100nM in combination with 5ng cDNA and Fast SYBR Green Mastermix (Invitrogen, #4385612). qPCR data were analysed using 2^-ΔΔCT^ method, and glyceraldehyde 3-phosphate dehydrogenase (GAPDH) was used as the reference gene.

### Myography

Thoracic aorta from mice aged 18-22 weeks were excised and placed immediately in ice-cold physiological salt solution (PSS; 130mM NaCl, 4.7mM KCl, 1.2mM MgSO_4_.7H_2_O, 24.9mM NaHCO_3_, 1.2mM KH_2_PO_4_, 2.5mM CaCl_2_ and 11.1mM glucose). Aortic segments were dissected to remove PVAT and cut into 2mm rings, as per standard myography protocol. Where there was a requirement for PVAT to be retained, aortic segments were dissected with minimal PVAT disruption (+PVAT) and compared to segments where PVAT was removed (-PVAT).

Aortic rings were mounted on 40μm wire in water baths (Danish Myo Technology) containing PSS, gassed with 95% O_2_ and 5% CO_2_. Vessels were normalised to 9.81mN, followed by exposure to KPSS (PSS with 62.5mM KCl) to establish maximal vascular contraction. Endothelial function was assessed by examining the acetylcholine relaxation response (1×10^−6^ M) following phenylephrine pre-contraction (1×10^−6^). Experiments with intact endothelial had >60% function preserved while denuded vessels had <20% function preserved. Following the initial normalisation, KPSS and endothelial function tests all experiments, were performed in modified PSS with 1mM CaCl_2_.

Cumulative concentration-response curves conducted in series for the α_1_-adrenoreceptor agonist phenylephrine (1×10^−9^M - 3×10^−5^M), the muscarinic (M)_2_-receptor agonist acetylcholine (ACh; 1×10^−9^M - 3×10^−5^M) and the nitric oxide donor sodium nitroprusside (SNP; 1×10^−10^M - 3×10^−6^M). Myograph readouts were generated using PowerLab hardware (ADInstruments, UK) and recorded (LabChart: v8). Phenylephrine vascular responses are expressed as mean contraction relative to KPSS contraction (%KPSS) ±SEM, while ACh and SNP responses are given as mean relaxation relative to phenylephrine pre-constriction (1×10^−6^M Phenylephrine) ±SEM.

### Adipose Transfer Studies

To examine the effect of secreted factors on vascular reactivity gonadal adipose transfer studies were performed. Paired female CaSR^fl/fl^ and CaSR^Ad-/-^ mice aged 18-21 weeks were sacrificed and gonadal adipose tissue dissected and placed immediately in ice-cold PBS. Aortas were prepared for myography studies as previously described. Isolated CaSR^fl/fl^ and CaSR^Ad-/-^ gonadal tissue was dissected to a final weight of 600mg and incubated at 37°C in 5mL reduced CaCl_2_ (1mM) PSS gassed with 95% O_2_ and 5% CO_2_ for 4h. Gonadal adipose tissue was halved by weight and transferred along with 2mL of adipose-conditioned media and 3mL reduced CaCl_2_ PSS to water baths containing CaSR^fl/fl^ or CaSR^Ad-/-^ aortas. Vessel segments were equilibrated for 30min before performing phenylephrine cumulative concentration-response curves (PE).

### Statistical Analysis

Data were analysed using GraphPad Prism (V9) and initially tested for normality and outliers. As appropriate either Student’s *t*-tests or Mann-Whitney test were used with two samples, one-way ANOVA or Kruskal-Wallis test were performed where groups >2, with post-hoc tests as appropriate. Data with multiple variables were analysed using either non-linear regression followed by an extra sum of squares F-test or two-way ANOVA followed by multiple comparison tests. Data are expressed as the arithmetic mean ± SEM for linear data and the geometric mean ± geometric standard deviation for logarithmic data (Rt-qPCR results).

## Results

We generated an adipocyte-specific CaSR knockout mouse by crossing the CaSR floxed mouse line (CaSR^fl/fl^) to mice possessing the AdipoQ-Cre BAC transgene to ensure CaSR expression was abolished in mature adipocytes only (CaSR^Ad-/-^) (Supplementary figure 1a). Successful excision of CaSR exon 3 was established in adipocytes that co-expressed Cre recombinase, with no corresponding band seen in non-adipocyte tissue (Supplementary Figure 1b). A reduction in CaSR exon 3 mRNA expression was confirmed in primary adipocytes isolated from the gonadal fat pads of a mixed population of both male and female CaSR^fl/fl^ and CaSR^Ad-/-^ mice (Supplementary Figure 1c).

Following confirmation of CaSR deletion, we assessed body weight in both sexes from 5 to 18 weeks of age. Female CaSR^Ad-/-^ mice showed a small but consistent reduction in body mass when compared to littermate controls (Figure 1a). In contrast, male body weight was unaffected by CaSR deletion (Figure 1b). We went on to determine whether this reduction in body mass was due to a reduction in adiposity. In females, both inguinal and gonadal fat pads were slightly smaller. However, this was not statistically significant suggesting that if CaSR deletion affects adiposity, it is a cumulative effect of all depots rather than a reduction in any specific adipose depot (Figure 1c). Male adipose depot size was unaffected by CaSR deletion (Figure 1d). Tibia length was unchanged in both sexes suggesting that female CaSR^Ad-/-^ mice are not simply smaller than CaSR^fl/fl^ littermates (figure 1e and f).

**Figure 1:**
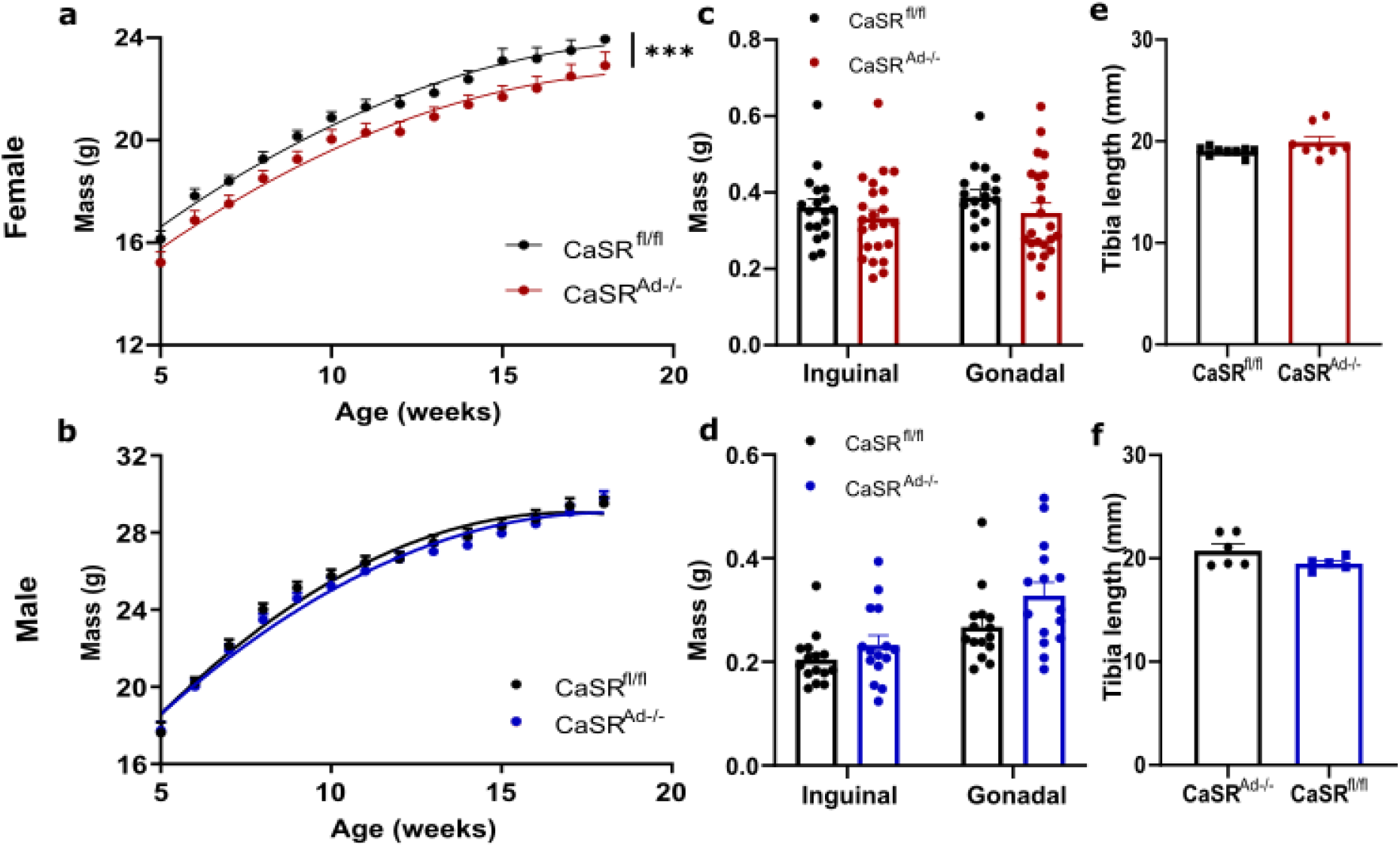
Adipocyte-specific CaSR deletion reduces female body mass. (a-b) Growth curves from 5-18 weeks of age in CaSR^fl/fl^ and CaSR^Ad-/-^ mice. Female (a) CaSR^Ad-/-^ were significantly smaller while males (b) were unaffected. Inguinal and gonadal fat pad mass in female (c) and male (d) CaSR^fl/fl^ and CaSR^Ad-/-^ were comparable at 18 weeks old. Tibia length was unaffected by CaSR deletion in both female (e) and male (f) mice. Data are presented as mean ± SEM for all graphs. Analysis by non-linear regression in (a) and (b), and students t-test in (c-f). ***P<0.001.

As female adipose depot size did not significantly change, we went on to investigate whether CaSR deletion influenced the adipocytes themselves. Histological analysis of gonadal adipose tissue demonstrated that on average wildtype female adipocytes were larger than male (Figure 2a). Following frequency distribution analysis, it became evident that CaSR deletion caused adipocytes in the female mice to shift leftwards becoming on average smaller than in the CaSR^fl/fl^ littermates (Figure 2a, b and d). Correspondingly, this led to an increase in the number of adipocytes per a region of interest (Figure 2e). In contrast, male adipocyte size and number were unaffected by CaSR deletion (Figure 2a, c, f and g). When female subcutaneous inguinal fat was analysed, we found no difference in adipocyte size between the genotypes suggesting that CaSR deletion is of greater significance in female visceral adipocyte function (Supplementary Figure 2a and b).

**Figure 2:**
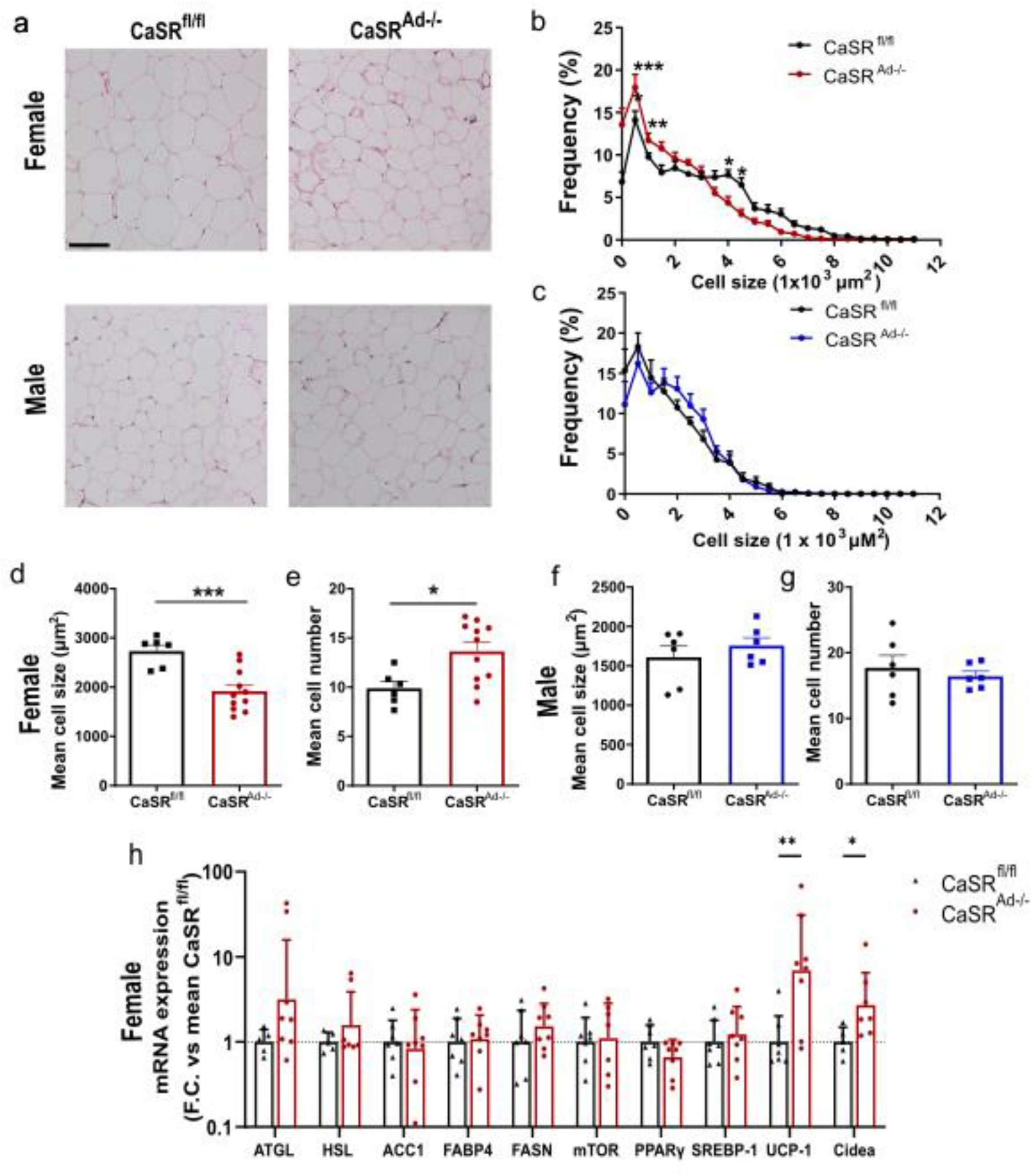
CaSR deletion reduces adipocyte size in female gonadal adipose tissue. (a) Representative images of gonadal adipose tissue from female and male CaSR^fl/fl^ and CaSR^Ad-/-^ mice. Size and frequency distribution of adipocytes from gonadal adipose depots in (b) female and (c) male CaSR^fl/fl^ and CaSR^Ad-/-^ mice. Mean adipocyte size in females (d) was reduced with a corresponding increase in mean cell number (e). Male adipocyte size (f) and number (g) were unaffected by CaSR deletion. (h) Rt-qPCR analysis of female primary adipocytes expressed as fold change (F.C) relative to the mean CaSR^fl/fl^ level of expression. ATGL: Adipose triglyceride lipase, HSL: Hormone-sensitive lipase, ACC1: Acetyl-CoA carboxylase, FABP4: Fatty acid binding protein 4, FASN: Fatty acid synthase, mTOR: mammalian target of rapamycin, PPARγ: Peroxisome proliferator-activated receptor gamma, SREBP-1: Sterol regulatory element-binding protein-1, UCP-1: Uncoupling protein 1, Cidea: Cell death-inducing DDFA like effector A. Data are presented as mean ± SEM in (a-g), and qPCR data is expressed as the geometric mean ± geometric standard deviation (h). Scale bar 100μm. Analysis by two-way ANOVA in (b) and (c), students t-test or Mann-Whitney test in (d-h) as determined by prior normality tests. * P<0.05, **P< 0.01, ***P<0.001.

To identify the mechanism underlying the reduction in female visceral adipocyte size in CaSR^Ad-/-^ compared to control mice we examined the expression of a panel of metabolic genes (Figure 2h). As CaSR activation has previously been described to be anti-lipolytic [8] and to suppress the expression of adipose triglyceride lipase (ATGL) and hormone-sensitive lipase (HSL), we first determined whether CaSR deletion altered the expression of these genes [6]. While ATGL did seem to increase in CaSR^Ad-/-^ gonadal adipocytes this did not reach statistical significance; HSL was comparable between genotypes suggesting that while CaSR activation may be suppress lipolysis, inhibition does not increase lipolytic activity. Therefore, we further investigated a panel of genes important for healthy adipocyte function. Most genes were unaffected by CaSR deletion; however, uncoupling protein 1 (UCP-1) and cell death inducing DFFA-like effector A (Cidea) were strongly upregulated in CaSR^Ad-/-^ adipose. Both Cidea and UCP-1 expression are markers of brown adipose tissue and are switched on when white adipocytes undergo ‘beiging’ [15]. Further work will be needed to understand the mechanisms whereby CaSR regulates these genes and whether gonadal adipocyte behaviour is altered by the increased expression of brown fat markers.

To understand whether these molecular and morphological changes had any systemic effects we investigated circulating concentrations of both triglycerides and free fatty acids (FFA). In female CaSR^Ad-/-^ mice triglyceride levels were significantly reduced compared to CaSR^fl/fl^ mice; in contrast FFA concentrations were comparable between the two genotypes.

Multiple studies have established a functional role for CaSR within the vasculature including in endothelial and vascular smooth muscle cells (vSMCs) [16-18]. Although it has been shown that CaSR is involved in neuronal signalling within the perivascular adipose tissue (PVAT) it has yet to be shown whether CaSR is expressed in perivascular adipocytes themselves [19]. First, we established whether CaSR expression levels were comparable across gonadal, inguinal and perivascular fat depots in female C57Bl/6 mice. CaSR expression within gonadal and inguinal adipocytes was similar, however PVAT expression was significantly higher (Supplementary Figure 3a). When PVAT CaSR expression is compared between sexes female mice have significantly higher levels of CaSR expression (Supplementary figure 4b), suggesting CaSR activity may again be more significant in female adipose tissue than male. To investigate whether this is the case, we utilised our CaSR^Ad-/-^ mouse. Confirmation of CaSR deletion in PVAT was achieved through the use of primers neighbouring exon 3 (Supplementary Figure 1a) with successful amplification of the recombination product only occurring in Cre recombinase positive mice (Supplementary Figure 3c).

Following confirmation of the successful deletion of CaSR in perivascular adipocytes we utilised wire myography to determine the role adipocyte CaSR plays in vascular reactivity. Experiments with aortic rings were performed in modified physiological saline solution (PSS) containing 1mM Ca^2+^. This concentration has been successfully used to previously investigate CaSR activity in vSMCs [18].

In the absence of PVAT (PVAT -), CaSR adipocyte specific deletion had no effect on phenylephrine induced contraction nor on acetylcholine induced relaxation in female aortas (Figure 4a and b). However, CaSR^Ad-/-^ vessels were significantly more sensitive to sodium nitroprusside (SNP) suggesting that CaSR deletion within in adipocytes increases the sensitivity of vSMCs to nitric oxide (NO) (Figure 4c). Interestingly, in endothelial denuded vessels CaSR deletion significantly reduced female aortic response to phenylephrine and significantly increased sensitivity to SNP (Figure 4d and e). In contrast, CaSR deletion had no basal effect on male PVAT – aortic rings. (Supplementary figure 4a - e).

**Figure 3:**
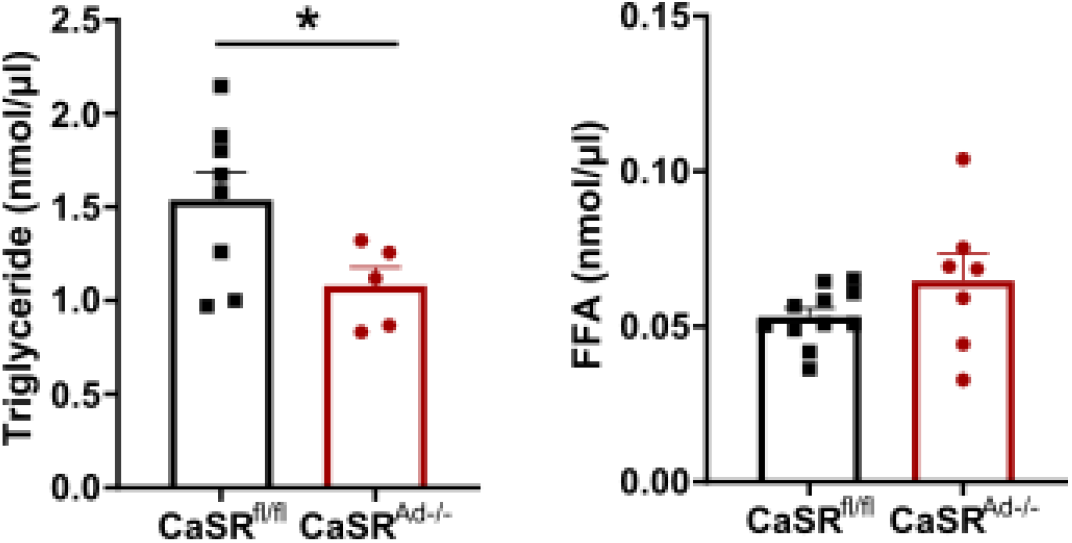
Plasma triglyceride concentrations are reduced in female CaSR^Ad-/-^ mice. Plasma(a) triglyceride and (b) free fatty acid (FFA) concentrations in female CaSR^fl/fl^ and CaSR^Ad-/-^ mice. Data are presented as mean ± SEM. Analysis by student t-test in (a) and (b). *P<0.05.

**Figure 4:**
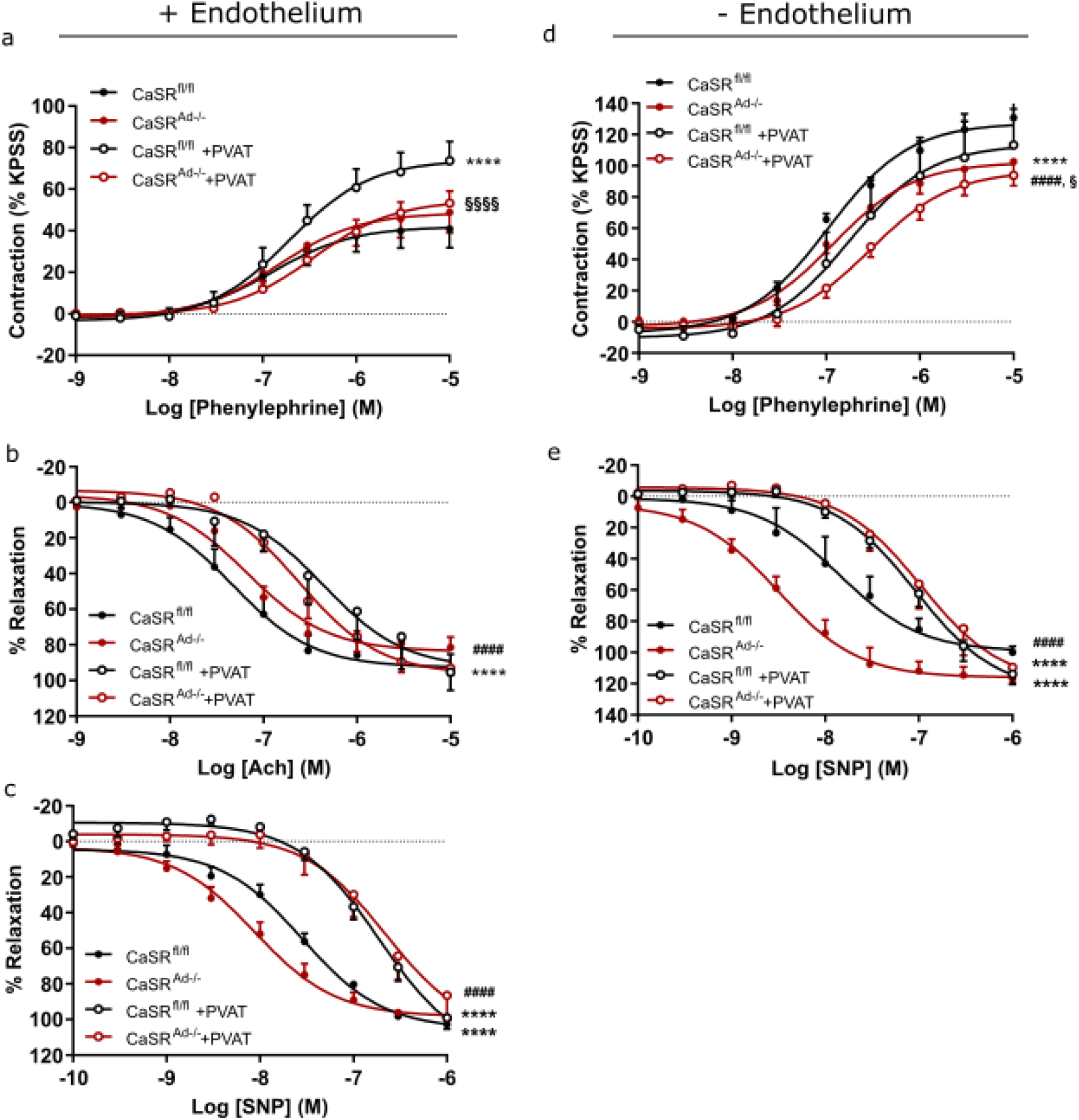
CaSR deletion in female PVAT reduces contraction to phenylephrine and increases sensitivity to SNP. Aortic rings from female CaSR^fl/fl^ and CaSR^Ad-/-^ were equilibrated in PSS containing 1mM Ca^2+^. Following equilibration vessels with intact endothelium were exposed to rising concentrations of (a) phenylephrine, (b) acetylcholine (Ach) and (c) sodium nitroprusside (SNP). Denuded vessels, with <20% endothelial function, were exposed to rising concentrations of (d) phenylephrine and (e) SNP. Data are expressed as mean ± SEM. Analysis was performed by non-linear regression and compared using an extra sum-of-squares F-test. ****P<0.0001 vs CaSR^fl/fl^ vessels; ####P<0.0001 vs CaSR^Ad-/-^ vessels; §P<0.05, §§§§P<0.0001 vs CaSR^fl/fl^ + PVAT.

PVAT intact (PVAT+) aortic rings from both male and female CaSR^fl/fl^ mice had increased contractile response to phenylephrine compared to PVAT -vessels (Figure 4a and Supplementary Figure 4a). Suggesting that in this mouse strain PVAT favours a pro-contractile environment.

As previously, male PVAT + CaSR^Ad-/-^ aortas were comparable to CaSR^fl/fl^ littermates (Supplementary Figure 4a). In contrast, PVAT+ vessels from female CaSR^Ad-/-^ mice were protected from the pro-contractile response of PVAT (Figure 4a). In both male and female vessels, PVAT reduced the sensitivity to acetylcholine and SNP, but CaSR deletion had no effect (Figure 4b and c; Supplementary Figure 4b and c). In endothelial denuded vessels PVAT significantly reduced contraction to phenylephrine in both CaSR^fl/fl^ and CaSR^Ad-/-^. Therefore, in female mice CaSR regulation of PVAT mediated contraction is likely to be dependent on the endothelium.

To determine the possible mechanisms through which CaSR regulates PVAT-mediated vascular reactivity in female mice, we probed the mRNA expression of a range of adipokines released by aortic perivascular adipose tissue. Adiponectin expression was significantly upregulated in CaSR^Ad-/-^ PVAT, while omentin expression was significantly downregulated (Figure 5a). Apelin, leptin, resistin and visfatin were all statistically unaffected by CaSR deletion. PVAT can also secrete factors that are well known for their endothelial-mediated vascular reactivity such as nitric oxide (NO), angiotensin II, endothelin-1 (EDN-1) and prostacyclin (PGI_2_). However, the expression of endothelial nitric oxide synthase (eNOS), angiotensin-converting enzyme (ACE), EDN-1 and prostaglandin-1 synthase (PTGIS) were statistically unchanged by CaSR deletion (Figure 5b).

**Figure 5:**
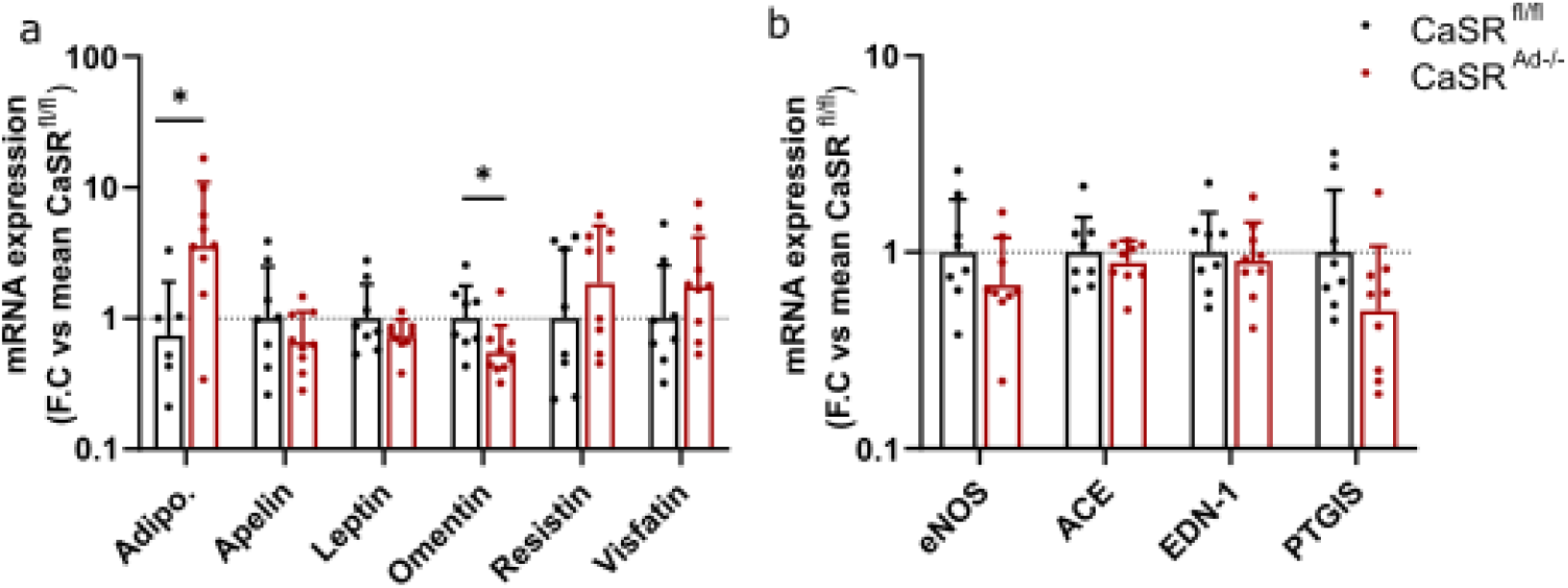
CaSR deletion increases adiponectin expression in female perivascular adipose tissue. Rt-qPCR analysis of (a) adipokine and (b) vascular reactive factor expression in perivascular adipose tissue from female CaSR^fl/fl^ and CaSR^Ad-/-^ mice. Data are presented as fold change (F.C.) relative to the mean CaSR^fl/fl^ level of expression. Adipo: Adiponectin, eNOS: endothelial nitric oxide synthase, ACE: angiotensin 1 converting enzyme, EDN-1: Endothelin I, PTGIS: Prostaglandin-1 synthase. Data are presented as the geometric mean ± geometric standard deviation. Analysis was done by students t-test or Mann-Whitney test as determined by prior normality tests. * P<0.05.

Adiponectin is a known vasodilator having both paracrine effects when secreted from PVAT but also participating in endocrine signalling when secreted from visceral adipose tissue. To explore whether adiponectin secretion is the protective mechanism through which female CaSR^Ad-/-^ aortas respond less to phenylephrine we incubated aortic rings with either their own gonadal adipose tissue or from the alternate genotype. As expected CaSR^fl/fl^ aortas exposed to CaSR^fl/fl^ gonadal fat demonstrated exaggerated contraction in response to phenylephrine, confirming that vessels were responding to a secreted factor (Figure 6a). Surprisingly, when CaSR^fl/fl^ aortas were incubated with CaSR^Ad-/-^ adipose tissue the vessel again exhibited heightened contractility. In contrast, when CaSR^Ad-/-^ vessels were incubated with CaSR^Ad-/-^ adipose tissue they contracted to the same degree as in the absence of fat confirming the findings in Figure 4a. Next, CaSR^Ad-/-^ vessels were incubated with adipose tissue from CaSR^fl/fl^ mice. Surprisingly, the CaSR^Ad-/-^ vessel only contracted to the same degree as vessels in the absence of adipose tissue (Figure 6b). Taken together these data suggest that although adiponectin expression is altered in CaSR^Ad-/-^ PVAT, the protective effect of adipocyte specific CaSR deletion is driven by chronic changes in the vessel itself.

**Figure 6:**
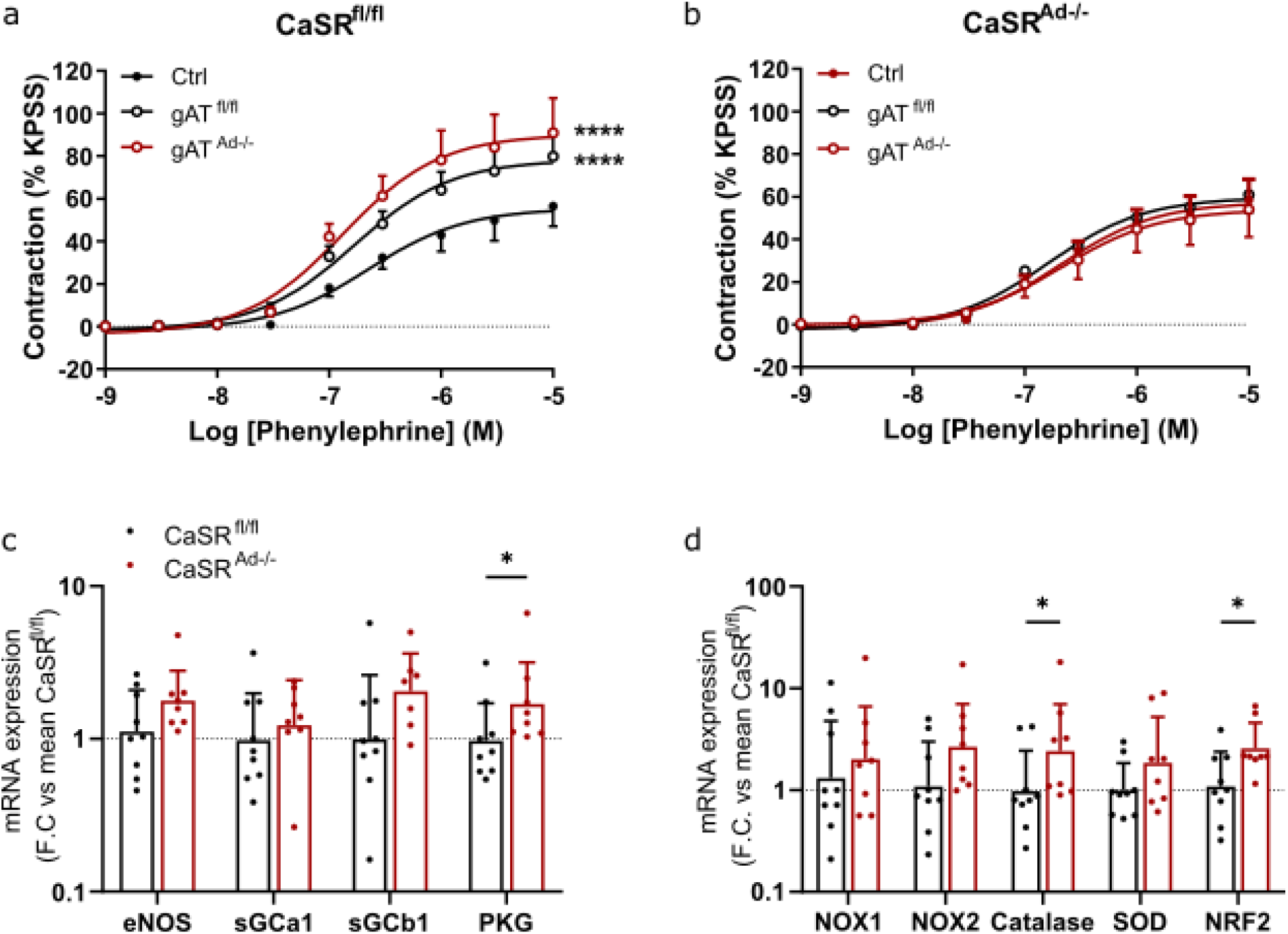
Female CaSR^Ad-/-^ aortas upregulate antioxidant signalling pathways to protect against adipose derived contractile factors. Aortas from (a) CaSR^fl/fl^ and (b) CaSR^Ad-/-^ were exposed to gonadal adipose tissue and pre-conditioned PSS from either their own or the alternate genotype. Vessels were then subjected to rising concentrations of phenylephrine. qPCR analysis of whole aorta with PVAT removed for mediators of (c) nitic oxide and (d) reactive oxygen species pathways. eNOS: endothelial nitric oxide synthase; sGCa1: soluble guanylyl cyclase A1; sGCb1: soluble guanylyl cyclase B1; PKG: protein kinase G; NOX1: NADPH oxidase 1; NOX2: NADPH oxidase 2; SOD: superoxide dismutase; NRF2: nuclear factor erythroid 2-related factor 2. Analysis in (a) and (b) was performed by non-linear regression and compared using an Extra sum-of-squares F-test. ****P<0.0001 vs CaSR^fl/fl^ vessels. Data in (c) and (d) are presented as the geometric mean ± geometric standard deviation. Analysis students t-test or Mann-Whitney test as determined by prior normality tests. * P<0.05.

To rule out structural changes to the vessel causing the observed phenotype, we undertook histological analysis of the aortas. No gross morphological changes were seen in the vessel wall of either male or female CaSR^Ad-/-^ vessels, suggesting that structural changes to the medial layer are not the cause of reduced contraction in female CaSR^Ad-/-^ vessels (Supplementary Figure 5).

To identify potential mechanisms that might underly the functional changes that we observed we ran qPCR experiments on whole aortas with the PVAT removed. Starting with the NO pathway we found no changes in the expression of eNOS nor soluble guanylyl cyclase (sGCa1 and sGCb1) but did find that protein kinase G (PKG) expression was significantly upregulated. This potentially explains why female aortas had heightened responsiveness to SNP (Figure 6c). Secondly, we examined mediators of reactive oxygen species signalling. While NADPH oxidase (NOX) 1 and 2 and superoxide dismutase (SOD) expression were unaffected by CaSR deletion, both catalase and nuclear factor erythroid 2-related factor 2 (NRF2) were significantly upregulated in the female aorta (Figure 6d). These observations suggest that vessels from female CaSR^Ad-/-^ mice might be protected from adipose induced contraction via an upregulation of antioxidant pathways.

## Discussion

A high calcium diet is generally considered to have beneficial effects in obesity and on blood pressure regulation [21, 22]. However, the mechanisms underlying this effect are manifold and currently not well understood. Calcium has long been recognised as a second messenger but was established as a primary signalling molecule following the identification of the calcium sensing receptor (CaSR) [2]. In this study, we aimed to increase our understanding of CaSR signalling and its effect on basal adipocyte function and vascular reactivity in a transgenic model of adipocyte specific CaSR deletion (CaSR^Ad-/-^).

In contrast to the general positive effects of calcium on obesity, CaSR activation has been shown to be anti-lipolytic [8]. Therefore, we sought to ascertain whether the absence of CaSR would lead to a reduction in fat mass and smaller adipocytes. Deletion of adipocyte CaSR led to a ~5% reduction in total body weight but surprisingly in female mice only. Although there was no significant effect of CaSR deletion on the size of specific adipose depots there was a profound shift in adipocyte size and number. Reduced adipocyte size and increased number (hyperplasia) are generally considered to correlate with improved metabolic function. Indeed, female CaSR^Ad-/-^ had significantly lower levels of plasma triglyceride. In contrast, male CaSR^Ad-/-^ mice were comparable to their wildtype counterparts. This may, in part, explain why Sundararaman *et. al*. [12] did not see any effect of CaSR deletion on a high fat diet as their use of a mixed sex population may have obscured the phenotype in female mice.

To determine the mechanism behind the reduction in adipocyte size in female mice we screened several important metabolic genes. Surprisingly, we found CaSR deletion had no effect on ATGL or HSL suggesting that CaSR inhibition does not necessarily generate the opposite effects of CaSR activation. Likewise, no change in lipogenic genes including PPARγ and FASN were observed. This is in keeping with our previous findings in 3T3-L1 cells where negative modulation of CaSR with NPS-2143 alone had no effect [7]. Interestingly, in this study we identified a strong upregulation of both UCP-1 and Cidea expression. UCP-1 and Cidea are considered brown adipose tissue markers [23], but can be upregulated in white adipose upon ‘beiging’ or ‘browning’. Normally, this process is associated with subcutaneous depots rather than visceral fat therefore further work will be necessary to understand the functional consequences of this change and how CaSR regulates UCP-1 and Cidea expression.

Perivascular adipose tissue (PVAT) is a significant regulator of vascular reactivity [24]. First reported to play an anti-contractile role in response to norepinephrine [25], PVAT has also been identified to secrete pro-contractile factors. In our studies aortic rings from male and female CaSR^fl/fl^ mice showed increased contraction to phenylephrine both when PVAT remained intact, and when incubated with gonadal adipose-conditioned media. These observations suggest that in this strain, a factor secreted from adipose tissue caused the increase in contraction. Interestingly, CaSR deletion reduced this contraction back to the PVAT negative baseline but only in female mice. Following adipose transfer studies with female tissue we were surprised to find that CaSR^Ad-/-^ gonadal fat seemed to behave normally when incubated with wildtype vessels. In contrast CaSR^Ad-/-^ vessels did not respond to either wildtype or CaSR^Ad-/-^ adipose tissue. We observed upregulation of antioxidant pathways in CaSR^Ad-/-^ vessels. Given that superoxide is thought to be one of the pre-dominant adipose-derived contractile factors [26, 27] it is possible that increased expression of catalase and NRF2 may negate any pro-contractile ROS production by adipose tissue. Further studies will be necessary to determine whether reactive oxygen species (ROS) production differs between the two mouse strains.

Understanding CaSR signalling in tissues beyond direct calcium homeostasis is important due to the large number of patients prescribed CaSR-positive allosteric modulators such as cinacalcet and the high frequency of CaSR polymorphisms. Here we demonstrate that CaSR is an important regulator of both visceral and perivascular adipose function exclusively in female mice. Female cardiovascular and metabolic health has often been overlooked and therefore it will be important to continue these studies to identify pathways regulated by CaSR and their role in human health and disease.

## Acknowledgements

Our thanks go to Lynn Stevenson at the University of Glasgow histology service.

## Author Contributions

Conceptualization, E.H, J.L and L.D; Methodology, E.H and L.D; Investigation, E.H, Z. G-G, J.M, and L.D; Formal Analysis, E.H, Z. G-G, and L.D; Writing-Original Draft, E.H. and L.D; Writing-Reviewing and editing, Z. G-G and J.L; Supervision, J.L and L.D; Funding Acquisition, E.H and J.L.

