## Supplementary Methods and Data for "The Calcium-sensing receptor is an important regulator of female visceral and perivascular adipocyte function"

**Supplementary Materials**

**Table 1: Genotyping primers to confirm the presence of cre recombinase, CaSR exon 3 floxed sites and CaSR exon 3 deletion.**

| Primer pair | Forward (5'>3') | Reverse (3'>5') |
|---|---|---|
| Cre Recombinase | GCCTGCATTACCGGTCGATGCAACGA | GTGGCAGATGGCGCGGCAACACCATT |
| CaSR floxed | TGTGTCAGTTTCATAGCCTGAAG | TGTCTATGGAAAGCCCAGA |
| CaSR Wildtype | CTTCAGAGGCCAGAGGTGTC | TGTCTATGGAAAGCCCAGA |
| CaSR Postneo | GACTTGCTATGTAGCCCAGAACTG | ACACCCCAAGTGCTCCTGATAACAG |

**Table 2: Primer pairs utilised for qPCR assessment of adipocyte and vascular function.**

| Primer Pair | Forward (5’>3’) | Reverse (3’>5’) |
|---|---|---|
| ACC1 | ATGGGCGGAATGGTCTCTTTC | TGGGGACCTTGCTTTCATCAT |
| ACE | CAGTTGCCCGGAATGAAACC | CTGGGCGGACTGGTAGATG |
| Adiponectin | CTTGTGCAGGTTGGATGGC | CAGCTCCTGTCATTCCAACA |
| Apelin | ATGAATGCTGAGGCTCTGCGT | TGCGGAACTTGATAAACAC |
| CaSR E2-E4 | ATGACTTCTGGTCCAATGAG | TGCGGAACTTGATAAACAC |
| CaSR | AGCACTGCGGCTCATGCTTTC | TCAGGGCCAGTGGTTGCTC |
| Catalase | ACATGGYCYGGGACYYCYGG | CAAGTTTTTGATGCCCTGGT |
| EDN-1 | TCGTGACTTTCCAAGGAGCT | CCAGGTGGCAGAAGTAGACA |
| FABP4 | TAAAAACACCGAGATTTCCTTCA | CCTTTCATAACACATTCCCACCA |
| FASN | TGCTCCCAGCTGCAGGC | GCCCGGTAGCTCTGGGTGTA |
| GAPDH | TGAACGGGAAGCTCACTGG | TCCACCACCCTGTTGCTGTA |
| Leptin | AAGAAGATCCCAGGGAGGAA | TGATGAGGGTTTTGGTGTCA |
| mTOR | GGCACATCTAGCAACGTGAG | CTGGTCATAGAAGCGAGTAG |
| NOX1 | TCCCTTTGCTTCCTTCTTGA | CCAGCCAGTGAGGAAGAGTC |
| NOX2 | CGCCCTTTGCCTCCATTCTC | CCTTTCCTGCATCTGGGTCTCC |
| NRF2 | TGAGCCAAGCTATAAGCCATGA | AATGGTTCTTGTGCCTGTGAA |
| Omentin | GTCTCTATTTCCTGCGCACG | CTCTGTTGCCTTGCTGACTG |
| Pgl2 | ATGCCTTGGAGTTTGGGAGA | TTCATCCTGGCCTTCTCCTC |
| Pkg | TGTCCTCGAAGAGACCCACT | GCAACGCTTTCTCTCCAAAC |
| PPARY | ATGCACTGCCTATGAGCACT | CAACTGTGGTAAAGGGCTTG |
| Resistin | TTCCTTGTCCCTGAACTGCT | TCTTCACGAATGTCCCACGA |
| Sgc1a | CCCCTGGTCAGGTTCCTAAG | GGAGACTCCCTTCTGCATTCT |
| SGC1B | TGCTGGTGATCCGCAATTATC | GGTTGAGGACTTTGCTTGCAG |
| SOD | GAGACCTGGGCAATGTGACT | TTGTTTCTCATGGACCACCA |
| UCP-1 | CGACTCAGTCCAAGAGTACTTCTCTTC | CGGGCTCAGAGTCACTACCACC |
| VISFATIN | GCAGAGCACAGTACCATAACG | TGGTGCCTCTGTACTTCTCG |

**Supplementary Data**

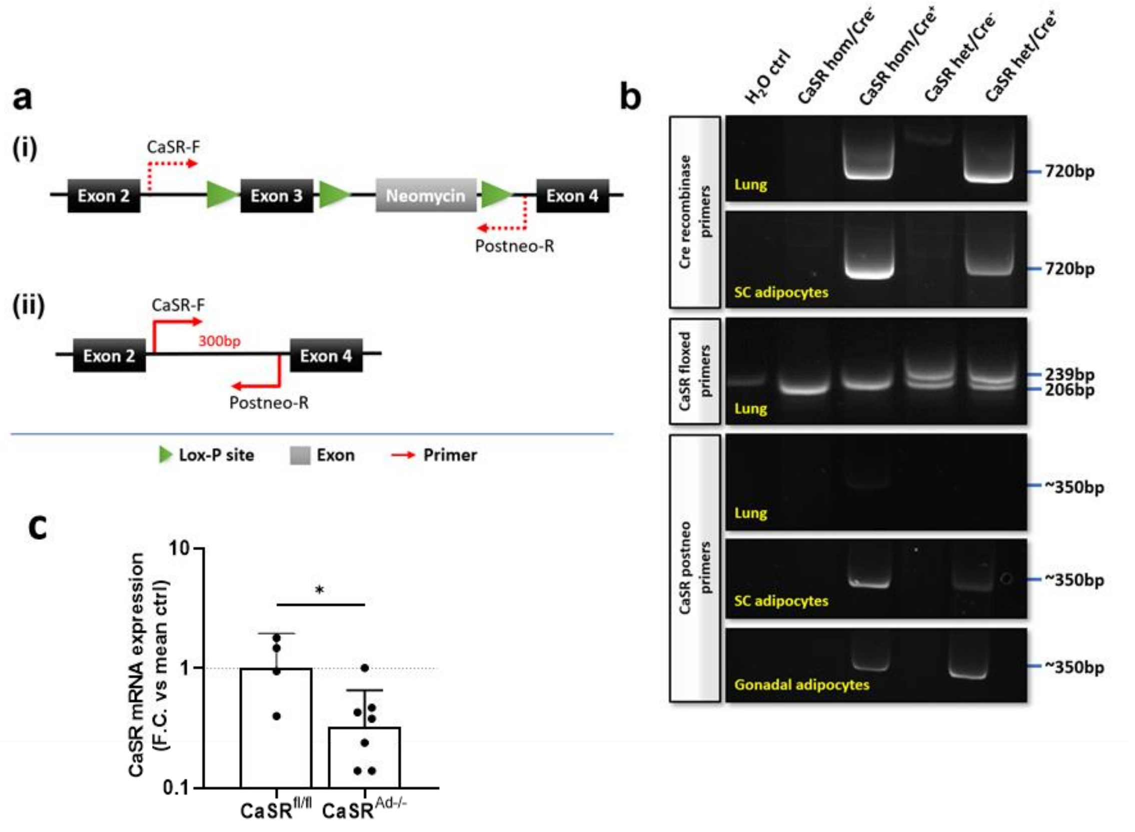


**Supplementary Figure 1: Verification of adipocyte specific deletion of CaSR.** (a) Illustrative diagram of the CaSR transgene designed by Toka et al showing the location of the CaSR postneo primer pair which only amplify following successful deletion of exon 3. (b) Representative End-point PCR gel demonstrating successful amplification of CaSR postneo product only occurs in adipocytes in the presence of Cre recombinase. (c) CaSR mRNA expression in significantly reduced in gonadal adipocytes isolated from $CaSR^{Ad-/-}$ mice. Data are presented as the geometric mean ± geometric standard deviation. Analysis by student t-test in (c). *P<0.05

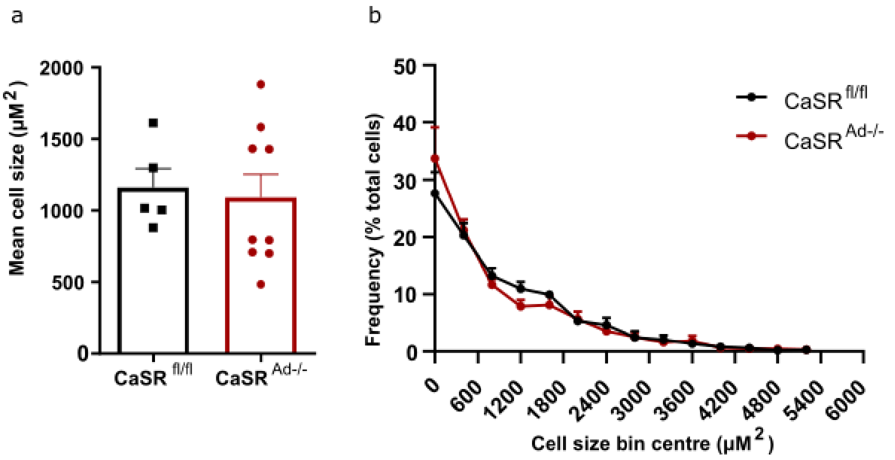


**Supplementary figure 2: Female subcutaneous inguinal adipocyte size is unaffected by CaSR deletion.** Mean size (a) and frequency distribution (b) of female inguinal adipocytes in $CaSR^{fl/fl}$ and $CaSR^{Ad-/-}$ mice. Data are presented as mean ± SEM.

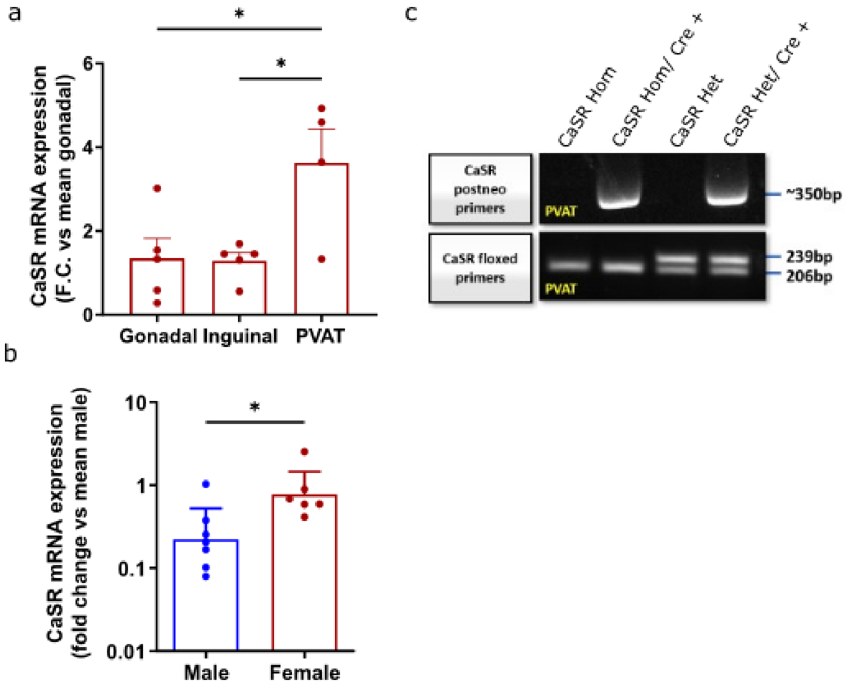


**Supplementary Figure 3: CaSR expression is significantly higher in female perivascular adipose tissue (PVAT) compared to male. (a)** CaSR mRNA expression is female C57Bl/6 gonadal, inguinal and PVAT . (b) CaSR mRNA expression in PVAT from male and female C57Bl/6 mice. (c) CaSR genotyping from PVAT. Amplification with CaSR postneo primer product only occurs in the presents of Cre recombinase. Data are presented as geometric means ± geometric standard deviation. Analysis by Kruskal-Wallis test in (a) and Mann-Whitney test in (b). *P<0.05.

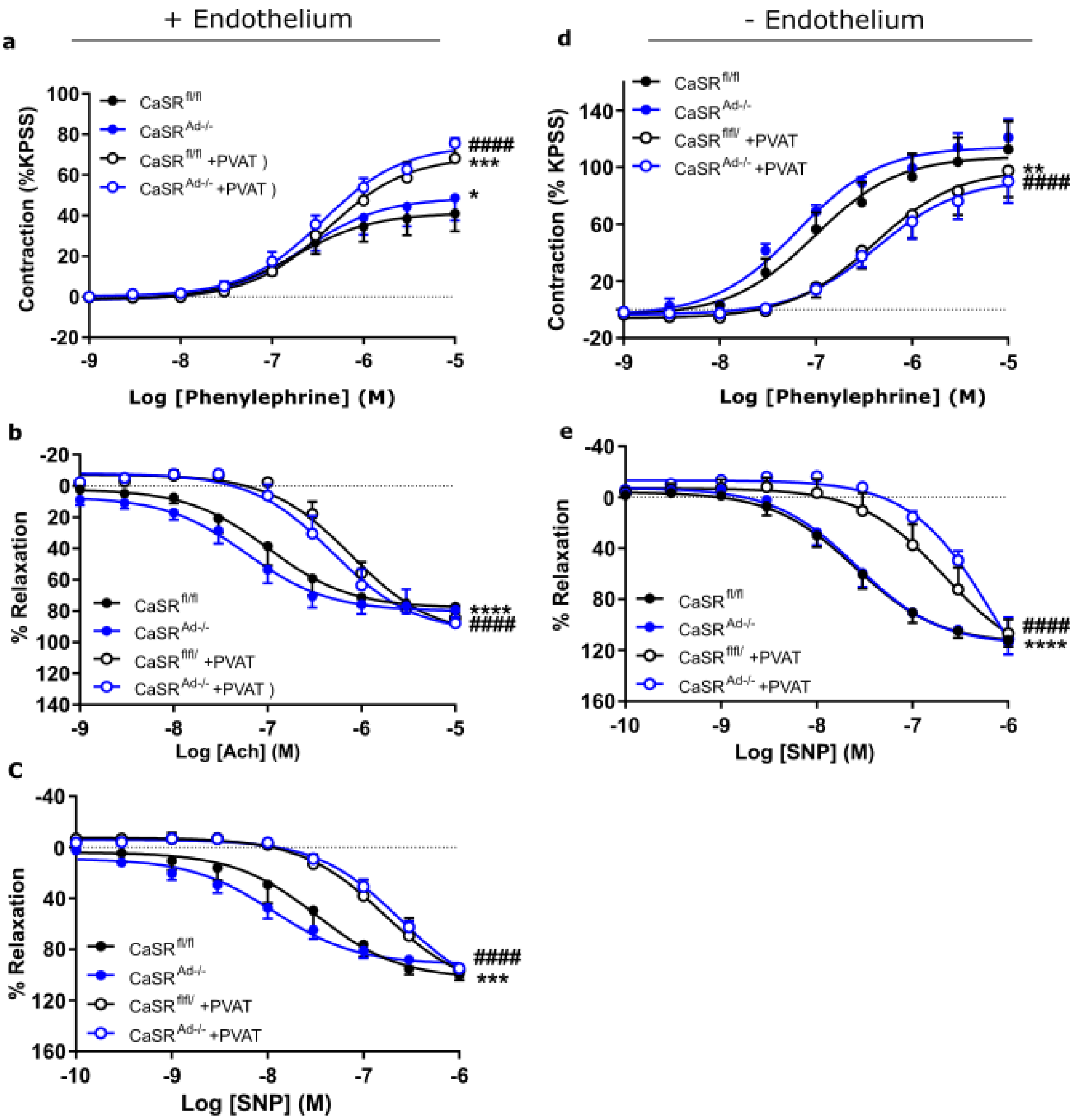


**Supplementary figure 4: CaSR deletion in perivascular adipose tissue has no effect on male vascular reactivity.** Aortic rings from male $CaSR^{fl/fl}$ and $CaSR^{Ad-/-}$ were equilibrated in PSS containing 1mM $Ca^{2+}$. Following equilibration vessels with intact endothelium were exposed to rising concentrations of (a) phenylephrine, (b) acetylcholine (Ach) and (c) sodium nitroprusside (SNP). Denuded vessels, with <20% endothelial function, were exposed to rising concentrations of (d) phenylephrine and (e) SNP. Data are expressed as mean ± SEM. Analysis was performed by non-linear regression and compared using an extra sum-of-squares F-test. ****P<0.0001, *** p<0.001, ** p<0.01 vs $CaSR^{fl/fl}$ vessels; ####P<0.0001 vs $CaSR^{Ad-/-}$ vessels.

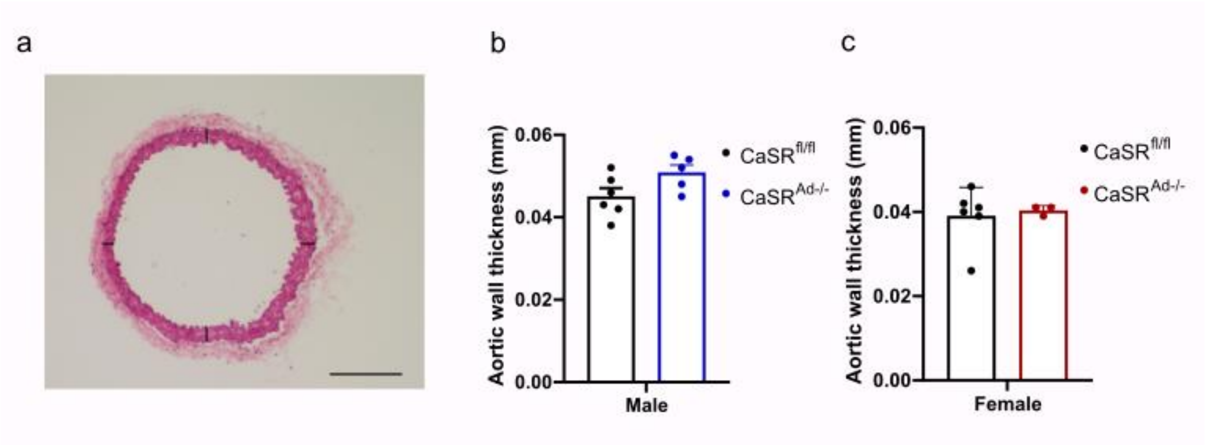


**Supplementary figure 5: Aortic wall thickness in male and female mice is unaffected by CaSR deletion.** (a) Representative image of cross section of mouse aortic wall. Black lines represent the four different locations where aortic thickness was measured (Scale bar represents 200um). No significant differences were observed between (b) male (p=0.072) and (c) female (p=0.754) $CaSR^{fl/fl}$ and $CaSR^{Ad-/-}$ mice. Data are presented as mean ± SEM. Analysis was performed by unpaired-t test.
